# Phylogeny-Informed Random Forests for Human Microbiome Studies

**DOI:** 10.1101/2025.11.04.686494

**Authors:** Hyunwook Koh

## Abstract

Random Forest is a widely used tree-based ensemble learning algorithm that efficiently captures complex nonlinear relationships and higher-order feature interactions with no distributional assumptions to be satisfied. It is also well-suited to human microbiome studies, where the data are highly skewed, overdispersed, discrete, and irregular. Here, I pay particular attention to the phylogenetic tree information that reflects evolutionary ancestry and functional relatedness among microbial features. Proper incorporation of phylogenetic tree information into microbiome data analysis has provided new insights and improved analytical performance. In this paper, I introduce an extension of the Random Forest algorithm that incorporates phylogenetic tree information, named Phylogeny-Informed Random Forests (PIRF), to improve predictive accuracy in human microbiome studies. The core mechanism of PIRF lies in its localized approach; rather than treating all features as competing globally to be selected or weighted, PIRF identifies informative features within each phylogenetic cluster (i.e., a localized group of microbial features that are evolutionarily and functionally related), thereby enriching functional representations while reducing tree correlation. I demonstrate the high predictive accuracy of PIRF, compared with other off-the-shelf tools, across seven benchmark tasks: four classification problems (gingival inflammation, immunotherapy response, type 1 diabetes, and obesity) and three regression problems (cytokine level, age based on oral microbiome, and age based on gut microbiome).

**Importance:** PIRF is an extension of the Random Forest algorithm that incorporates phylogenetic tree information to improve predictive accuracy in human microbiome studies. PIRF can serve as a useful tool for microbiome-based disease diagnostics and personalized medicine. The software and tutorials are freely available as an R package, named `PIRF`, at https://github.com/hk1785/PIRF.

## 1 Introduction

Decision Tree [1] is a nonparametric learning algorithm that recursively partitions the feature space into multiple regions and assigns a predicted output to each region. Its hierarchical structure and the discrete nature of the resulting regions make it robust to complex nonlinear relationships and higher-order feature interactions. As a nonparametric method, it does not also require any distributional assumptions to be satisfied; as such, it is well-suited to highly skewed or irregularly distributed data.

In recent years, ensemble learning algorithms based on decision trees, such as Random Forest [2] and Gradient Boosting Machine [3], have gained increasing popularity. First, Random Forest [2] is a bootstrap aggregation method to combine predicted outputs from an ensemble of bagged decision trees, each decision tree built using a random subset of features. Of importance is here that, through random feature selection in the bootstrap aggregation process, Random Forest [2] decorrelates decision trees, thereby accelerating variance reduction and substantially improving predictive accuracy. On the other hand, Gradient Boosting Machine [3], a close competitor to Random Forest [2], refines a predictive model sequentially by fitting weak learners, typically shallow decision trees, to the residuals of prior models. Gradient Boosting Machine [3] shares many advantages of Random Forest [2], yet it learns slowly based on a small learning rate with additional hyperparameters (e.g., maximum tree depth and the number of boosting iterations) to be carefully tuned. Therefore, Random Forest [2] is often faster to train and more automatic in practice.

Random Forest [2] is well-suited to human microbiome studies to predict a host’s health or disease status based on the composition of microbial features (e.g., operational taxonomic units, amplicon sequence variants) in the human body. This is because of the high complexity of microbiome data; the data are highly skewed, zero-inflated, and overdispersed. Here, I pay particular attention to the fact that in human microbiome studies, a phylogenetic tree is often provided as supplementary material alongside microbial feature abundance data. It is constructed based on genetic sequence similarities among microbial features, and represents evolutionary ancestry and functional relatedness among microbial features [4]. Proper incorporation of phylogenetic tree into microbiome data analysis has provided new insights and improved analytic performance in *α*- and *β*-diversity estimation [5, 6], significance testing [7, 8], causal inference [9, 10], and predictive modeling [11, 12, 13, 14].

In this paper, I introduce an extension of the Random Forest [2] algorithm that incorporates phylogenetic tree information, named Phylogeny-Informed Random Forests (PIRF), to improve predictive accuracy in human microbiome studies. Motivating prior extensions of Random Forest are feature elimination [15, 16] and feature weighting [17] approaches designed to address high-dimensional settings by leveraging feature importance scores (e.g., increase in training loss when each feature is permuted or decrease in training loss when each feature is used for splitting). More specifically, feature importance scores have been used either to eliminate less informative features (i.e., to exclude them from the random feature selection process) [15, 16] or to assign higher weights to more informative features (i.e., to increase their likelihood of being included in the random subset of features) [17]. These approaches are intuitive and appealing for enhancing the strength of the fitted tree [2], and have demonstrated improved predictive accuracy in several empirical settings [15, 17, 16]. However, they also entail a potential drawback; that is, when certain features are selected more frequently (i.e., not equally likely), similar features tend to be used across decision trees, leading to an increase in correlation among decision trees. Such an inflated correlation undermines the variance-reduction benefit in the bootstrap aggregation process, ultimately reducing predictive accuracy. As a remedy, PIRF adopts a localized approach; instead of treating all features as competing globally to be selected or weighted, PIRF identifies informative features within each phylogenetic cluster (i.e., a localized group of microbial features that are evolutionarily and functionally related). For this, PIRF first partitions the microbial feature space into multiple phylogenetic clusters using phylogenetic tree information. PIRF then computes feature importance scores within each cluster and converts them into cluster-specific probabilities. Finally, the cluster-specific probabilities are integrated over all phylogenetic clusters to derive community-level selection probabilities. This localized strategy diversifies functional representations while mitigating tree correlation.

I demonstrate the high predictive accuracy of PIRF, compared with other off-the-shelf tools, using (i) four classification tasks on gingival inflammation [18], immunotherapy response [19], type 1 diabetes (T1D) [20], and obesity [21]; and (ii) three regression tasks on cytokine level [18], age based on oral microbiome [18], and age based on gut microbiome [21]. The software and tutorials are freely available as an R package, named `PIRF`, at https://github.com/hk1785/PIRF. Its core routines are built on the R package `ranger` [22], which is implemented in C++ and supports multi-core parallel computation. Its trained model can also be easily stored, shared, and applied to new observations for rapid prediction without the need for retraining.

## 2 Materials and Methods

### 2.1 Phylogenetic Clusters

Microbial features (e.g., operational taxonomic units, amplicon sequence variants) are taxonomic units defined and classified based on genetic sequence similarities, typically using 16S ribosomal RNA genes in amplicon sequencing or genome-wide sequences in shotgun metagenomics [4]. In the context of predictive modeling, they are referred as features or input variables, and represented by their measured quantities (typically, relative abundances), denoted as in Eq. (1).

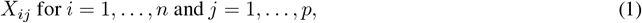

where *n* is the number of subjects and *p* is the number of features, possibly with the high dimensionality of *p* ≫ *n*.

To characterize the evolutionary and functional relatedness among microbial features, a phylogenetic tree is constructed based on their genetic sequence similarities [4]. It consists of leaves (representing microbial features), internal nodes (representing common ancestors), and branches that connect nodes. The length of each branch reflects the evolutionary distance (i.e., the degree of genetic divergence) between nodes. Microbial features with high genetic sequence similarity are considered phylogenetically related, sharing a recent common ancestor and exhibiting similar genetic characteristics and biological functions.

Phylogenetic clusters are localized groups of microbial features that are phylogenetically related. To identify these clusters, PIRF computes pairwise cophenetic distances between microbial features using the phylogenetic tree information [23], and then applies the partitioning around medoids algorithm [24] to partition *p* features into *k* clusters. Here, the cophenetic distance is a phylogenetic distance measure defined as the total branch length connecting two features to their most recent common ancestor (i.e., the nearest internal node) [23]. In microbiome data, most microbial features are rare variants with excessive zeros, which makes abundance-based distances ineffective for separating them into biologically meaningful clusters; as such, the cophenetic distance [23] has been widely used for clustering microbial features into distinct functional groups [8, 25].

The number of clusters (*k*) depends on how the phylogenetic relatedness is defined: (i) the most conservative definition yields fully disjoint clusters (*k* = *p*; each feature forms its own cluster), (ii) the most lenient definition yields a single cluster (*k* = 1; all features in one cluster); and (iii) a moderate definition yields 1 *< k < p*. For an optimal clustering resolution, PIRF identifies the number of clusters, denoted as *k*_*phylo*_, by maximizing the average silhouette width (i.e., a measure of how well each feature fits within its own cluster relative to others) [26] over the range of 2-10 clusters. Note that the two boundary cases, a single cluster (*k* = 1) and fully disjoint clusters (*k* = *p*), are non-phylogenetic clusters, whereas *k*_*phylo*_ clusters identified by PIRF are phylogenetic clusters derived from the phylogenetic tree. I denote *k*_*phylo*_ phylogenetic clusters as in Eq. (2).

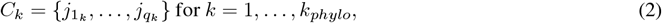

where *q*_*k*_ is the number of microbial features in the *k*-th cluster. These clusters are mutually exclusive and exhaustive: 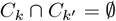 for *k* ≠ *k*′ and 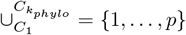.

### 2.2 Localized Feature Selection and Weighting

Feature importance scores quantify the contribution of each feature to predictive performance. Two notable approaches are: (i) permutation-based importance, which measures the increase in training loss when each feature is permuted (randomly shuffled), averaged over multiple permutations; and (ii) split-based importance, which measures the decrease in training loss from splits involving each feature, averaged over all bagged trees [27, 28]. PIRF adopts permutation-based importance because it is model-agnostic, enabling consistent comparison across different algorithms, less biased to continuous or high-cardinality categorical features, and captures feature interactions more effectively [27, 28].

PIRF computes feature importance scores within each phylogenetic cluster, denoted as in Eq. (3).

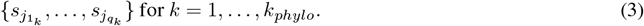

A negative feature score indicates that when a feature is permuted (randomly shuffled), predictive performance is improved, indicating no evidence of importance; hence, in Eq. (3), negative scores are set to zero for feature elimination, while positive scores are retained for feature selection. Then, the cluster-specific importance scores in Eq. (3) are converted into cluster-specific probabilities as in Eq. (4).

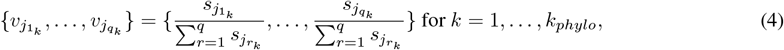

where 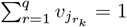. Finally, the cluster-specific probabilities in Eq. (4) are integrated over all phylogenetic clusters to derive community-level selection probabilities as in Eq. (5).

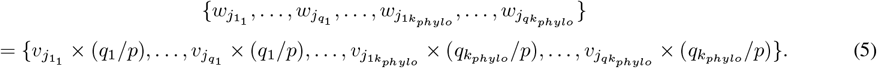

Here, each cluster-specific probability is multiplied by the ratio of its cluster size to the total number of features (i.e., if a feature belongs to the *k*-th cluster, its cluster-specific probability is multiplied by (*q*_*k*_*/p*)). This operation ensures (i) 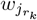 ‘s for *r* = 1, …, *q* and *k* = 1, …, *k*_*phylo*_ to form *p* probabilities for *p* features, satisfying the unit-sum constraint of 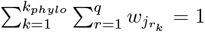; and (ii) every feature to have an equal opportunity of being weighted, regardless of its cluster size. The probabilities in Eq. (5) are feature-specific weights that quantify the relative importance of microbial features within their localized phylogenetic cluster. These probabilities provide a probabilistic mechanism for selecting subsets of features. That is, in contrast to the standard Random Forest [2], which selects features uniformly at random, PIRF selects features according to the probabilities in Eq. (5), prioritizing more informative microbial features.

Note that when the localized feature selection and weighting procedures from Eq. (3) to Eq. (5) are applied, the two non-phylogenetic clustering cases, a single cluster (*k* = 1) and fully disjoint clusters (*k* = *p*), correspond, respectively, to the one-cluster-based feature selection and weighting approach [17] and the standard Random Forest [2].

### 2.3 Other Technical Details

In PIRF, the number of randomly selected features can be tuned at both the cluster and global levels. Within each phylogenetic cluster containing *q*_*k*_ microbial features (*k* = 1, …, *k*_*phylo*_), I computed cluster-specific feature importance scores in Eq. (3) by averaging the ones from three candidate values: 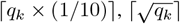, and ⌈log_2_ *q*_*k*_⌉. At the global level, I considered three analogous candidate values: ⌈*p*^*′*^ × (1*/*10)⌉, 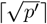, and ⌈log_2_ *p*^*′*^⌉, where *p*^*′*^ is the number of non-zero probabilities in Eq. (5), and selected the optimal value by minimizing out-of-bag error rates over 10,000 bagged decision trees.

I set all other hyperparameters to the default settings in the `ranger` package [22]: the minimum node size was 1 for classification and 5 for regression, with node splitting criteria based on Gini impurity for classification and variance reduction for regression (https://cran.r-project.org/web/packages/ranger/index.html). Additional user options are available in the `PIRF` software package (https://github.com/hk1785/PIRF).

## 3 Results

I evaluated the predictive accuracy of PIRF, relative to other off-the-shelf tools, as follows. For classification tasks, test error rate and test area under the curve (AUC) were employed, while for regression tasks, test root mean squared error (RMSE) and test mean absolute error (MAE) were employed, To estimate these predictive performance metrics, I partitioned the entire dataset into five folds: with four folds used for training and the remaining one fold used for testing. This process was repeated five times so that each fold was used once for testing, and the final estimates were averaged over the five iterations.

Details of the microbiome datasets I used and the other off-the-shelf tools I compared are provided in the following sections.

### 3.1 Microbiome Datasets

I considered seven benchmark tasks: (i) four classification problems on gingival inflammation [18], immunotherapy response [19], T1D [20], and obesity [21]; and (ii) three regression problems on cytokine level [18], age based on oral microbiome [18], and age based on gut microbiome [21]. For all these tasks, quality-control filters were applied uniformly, retaining only the subjects with more than 2,000 total read count and the microbial features with mean relative abundance greater than 0.00001. Additional preprocessing procedures and the resulting descriptive statistics are as follows.

- **Inflammation [18]:** For the gingival inflammation classification task, I used oral microbiome data from the salivary niche of participants in Baltimore, MD [18]. The dataset included 217 subjects (115 without inflammation; 102 with gingival inflammation) and 2,319 microbial features. The variation due to sample collection times and e-cigarette use was adjusted for using ConQuR [29].
- **Immunotherapy [19]:** For the immunotherapy response classification task, I used gut microbiome data on the efficacy of cancer immunotherapy in metastatic melanoma patients, compiled in the meta-analysis of [19]. The dataset included 257 subjects (168 non-respondents; 89 respondents) and 986 microbial features, collected from five different studies [30, 31, 32, 33, 34]. The study-specific variation was adjusted for using ConQuR [29].
- **T1D [20]:** For the T1D classification task, I used gut microbiome data from the non-obese diabetic mouse model experiment of [20]. The dataset included 521 subjects (167 without T1D; 354 with T1D) and 307 microbial features. The variation due to sample collection times and antibiotic use was adjusted for using ConQuR [29].
- **Obesity [21]:** For the obesity classification task, I used gut microbiome data from US-born residents in the American Gut Project (AGP) [21]. The dataset included 2,162 subjects (1,813 normal; 349 obese) and 2,162 microbial features.
- **Cytokine [18]:** For the cytokine level regression task, I used oral microbiome data from the salivary niche of participants in Baltimore, MD [18]. Cytokines are small signaling proteins secreted by immune and epithelial cells, playing essential roles in inflammation, immune responses, and intercellular communication. In this analysis, interleukin-8 (IL-8), a prototypical pro-inflammatory cytokine of the chemokine family, was used as a surrogate marker of cytokine level. The dataset included 190 subjects (minimum IL-8: 6.66; Q1 IL-8: 237.02; median IL-8: 532.12; Q3 IL-8: 890.49; maximum IL-8: 2359.28) and 2,323 microbial features. The variation due to sample collection times and e-cigarette use was adjusted for using ConQuR [29].
- **Age (Oral) [18]:** For the age (oral) regression task, I used oral microbiome data from the salivary niche of participants in Baltimore, MD [18]. The dataset included 217 subjects (minimum age: 18; Q1 age: 20; medium age: 24; Q3 age: 27; maximum age: 34) and 2,319 microbial features. The variation due to sample collection times and e-cigarette use was adjusted for using ConQuR [29].
- **Age (Gut) [21]:** I used gut microbiome data from US-born residents in AGP [21]. The dataset included 2,723 subjects (minimum age: 20; Q1 age: 35; medium age: 46; Q3 age: 59; maximum age: 69) and 2,111 microbial features.

### 3.2 Other Off-the-Shelf Tools

I evaluated eight other off-the-shelf tools in predictive accuracy as follows.

- **Random Forest [2]:** Standard random forest corresponding to fully disjoint clusters (*k* = *p*). The same parameter settings and training procedures as in PIRF were applied.
- **Random Forest (One Cluster) [17]:** Random forest with one-cluster-based feature selection and weighting (*k* = 1). The same parameter settings and training procedures as in PIRF were applied.
- **Gradient Boosting Machine [3]:** Gradient boosting trained with a learning rate of 0.0001, a maximum of 50,000 iterations, and early stopping after 10 rounds. Candidate interaction depths of 1–5 were tuned using 5-fold cross-validation, with cross-entropy loss for classification tasks and mean squared error for regression tasks. All other parameters were set to the defaults of the `xgboost` package [35] (https://cran.r-project.org/web/packages/xgboost/index.html).
- **Ridge Regression [36]:** Ridge regression trained with the regularization parameter tuned using 5-fold cross-validation, with cross-entropy loss for classification tasks and mean squared error for regression tasks. All other parameters were set to the defaults of the `glmnet` package (https://cran.r-project.org/web/packages/glmnet/index.html).
- **Lasso [37]:** Lasso regression trained with the regularization parameter tuned using 5-fold cross-validation, with cross-entropy loss for classification tasks and mean squared error for regression tasks. All other parameters were set to the defaults of the `glmnet` package (https://cran.r-project.org/web/packages/glmnet/index.html).
- **Elastic Net [38]:** Elastic net regression trained with the regularization and mixing parameters tuned using 5-fold cross-validation, with cross-entropy loss for classification tasks and mean squared error for regression tasks. All other parameters were set to the defaults of the `glmnet` package (https://cran.r-project.org/web/packages/glmnet/index.html).
- **Deep Neural Network (Keras) [39, 40]:** Two-layer neural network trained with the Adam optimizer [41], a learning rate of 0.0001 and an early stopping patience of 50. Candidate numbers of neurons per layer were [*p* × 0.1], [*p* × 0.2], [*p* × 0.3], [*p* × 0.4], [*p* × 0.5]; candidate dropout rates were 0.1–0.5; and candidate batch sizes were [*n* × 0.05], [*n* × 0.1], [*n* × 0.2], [*n* × 0.5]. These hyperparameters were tuned using the 5-fold validation-set approach, with cross-entropy loss for classification tasks and mean squared error for regression tasks. All other parameters were set to the defaults of the `keras3` package (https://cran.r-project.org/web/packages/keras3/index.html).
- **Deep Neural Network (Torch) [39, 40, 42]:** Two-layer neural network trained with the Adam optimizer [41], a learning rate of 0.0001 and an early stopping patience of 50. Candidate numbers of neurons per layer were [*p* × 0.1], [*p* × 0.2], [*p* × 0.3], [*p* × 0.4], [*p* × 0.5]; candidate dropout rates were 0.1–0.5; and candidate batch sizes were [*n* × 0.05], [*n* × 0.1], [*n* × 0.2], [*n* × 0.5]. These hyperparameters were tuned using the 5-fold validation-set approach, with cross-entropy loss for classification tasks and mean squared error for regression tasks. All other parameters were set to the defaults of the `torch` package [42] (https://cran.r-project.org/web/packages/torch/index.html).

### 3.3 Predictive Performances

The predictive performances across different tasks and methods are organized as follows.

- Table 1 reports the test error rates and AUC values for the four classification tasks: Inflammation [18], Immunotherapy [19], T1D [20], and Obesity [21] using each of the nine tools: PIRF, Random Forest [2], Random Forest (One Cluster) [17], Gradient Boosting Machine [3], Ridge Regression [36], Lasso [37], Elastic Net [38], Deep Neural Network (Keras) [39, 40], and Deep Neural Network (Torch) [39, 40, 42].
- Table 2 reports the test RMSE and MAE values for the three regression tasks: Cytokine [18], Age (Oral) [18], and Age (Gut) [21] using each of the nine tools: PIRF, Random Forest [2], Random Forest (One Cluster) [17], Gradient Boosting Machine [3], Ridge Regression [36], Lasso [37], Elastic Net [38], Deep Neural Network (Keras) [39, 40], and Deep Neural Network (Torch) [39, 40, 42].

**Table 1:** Test error rates and AUC values for PIRF and other existing off-the-shelf tools for the classification tasks of Inflammation, Immunotherapy, T1D, and Obesity (Unit: %). * RF, RF (One Cluster), GBM, Ridge, Lasso, EN, DNN (Keras), and DNN (Torch) represent Random Forest, Random Forest (One Cluster), Random Forest (Euclidean), Gradient Boosting Machine, Ridge Regression, Lasso, Elastic Net, Deep Neural Network (Keras), and Deep Neural Network (Torch), respectively.

| | Inflammation<br>$n = 217$<br>$p = 2,319$ | Immunotherapy<br>$n = 257$<br>$p = 986$ | T1D<br>$n = 521$<br>$p = 307$ | Obesity<br>$n = 2,162$<br>$p = 2,162$ |
| --- | --- | --- | --- | --- |
| Test Error Rate |  |  |  |  |
| PIRF | 10.20 | 8.15 | 11.13 | 15.91 |
| RF | 11.57 | 8.93 | 13.42 | 16.18 |
| RF (One Cluster) | 11.57 | 8.55 | 11.32 | 16.42 |
| GBM | 15.26 | 9.72 | 14.96 | 16.60 |
| Ridge | 40.53 | 29.93 | 20.91 | 16.48 |
| Lasso | 11.62 | 13.53 | 23.79 | 16.63 |
| EN | 12.07 | 19.01 | 22.83 | 16.58 |
| DNN (Keras) | 42.99 | 34.66 | 32.04 | 17.14 |
| DNN (Torch) | 28.56 | 20.27 | 32.04 | 16.28 |
| Test AUC |  |  |  |  |
| PIRF | 96.84 | 94.20 | 94.95 | 74.05 |
| RF | 96.17 | 91.77 | 93.51 | 71.57 |
| RF (One Cluster) | 96.21 | 93.14 | 94.74 | 73.89 |
| GBM | 92.27 | 87.00 | 93.28 | 66.42 |
| Ridge | 79.49 | 87.31 | 79.61 | 70.98 |
| Lasso | 94.20 | 82.26 | 79.24 | 68.06 |
| EN | 94.92 | 75.75 | 80.08 | 72.01 |
| DNN (Keras) | 55.99 | 62.48 | 55.37 | 58.53 |
| DNN (Torch) | 79.87 | 85.38 | 55.81 | 69.12 |

**Table 2:** Test RMSE and MAE values for PIRF and other existing off-the-shelf tools for the regression tasks of Cytokine, Age (Oral), and Age (Gut). * RF, RF (One Cluster), GBM, Ridge, Lasso, EN, DNN (Keras), and DNN (Torch) represent Random Forest, Random Forest (One Cluster), Random Forest (Euclidean), Gradient Boosting Machine, Ridge Regression, Lasso, Elastic Net, Deep Neural Network (Keras), and Deep Neural Network (Torch), respectively.

| | Cytokine<br>$n = 190$<br>$p = 2,323$ | Age (Oral)<br>$n = 217$<br>$p = 2,319$ | Age (Gut)<br>$n = 2,723$<br>$p = 2,111$ |
| --- | --- | --- | --- |
| Test RMSE |  |  |  |
| PIRF | 241.20 | 2.20 | 11.53 |
| RF | 259.05 | 2.36 | 11.69 |
| RF (One Cluster) | 244.85 | 2.31 | 11.79 |
| GBM | 277.42 | 3.36 | 12.52 |
| Ridge | 523.76 | 5.21 | 14.19 |
| Lasso | 441.05 | 3.93 | 13.03 |
| EN | 404.24 | 5.20 | 13.34 |
| DNN (Keras) | 471.58 | 4.22 | 13.18 |
| DNN (Torch) | 452.75 | 4.15 | 13.29 |
| Test MAE |  |  |  |
| PIRF | 144.18 | 1.35 | 9.37 |
| RF | 169.81 | 1.56 | 9.50 |
| RF (One Cluster) | 147.29 | 1.47 | 9.65 |
| GBM | 182.50 | 2.60 | 10.26 |
| Ridge | 385.15 | 3.89 | 12.09 |
| Lasso | 363.90 | 3.30 | 10.09 |
| EN | 328.73 | 3.42 | 10.70 |
| DNN (Keras) | 368.98 | 3.51 | 11.00 |
| DNN (Torch) | 370.54 | 3.49 | 11.19 |

To summarize, PIRF showed the strongest predictive performance across all tasks: for classification, it achieved the lowest test error rate and the highest test AUC (Table 1); for regression, it achieved the lowest test RMSE and the lowest test MAE (Table 2).

Among the comparison methods, tree-based models - including Random Forest [2], Random Forest (One Cluster) [17], and Gradient Boosting Machine [35] - generally outperformed linear models - including Ridge Regression [36], Lasso [37], and Elastic Net [38] - for most tasks (Tables 1–2), which is likely due to their ability to capture nonlinear relationships and higher-order feature interactions. Note also that the recently popular Deep Neural Networks [39, 40] implemented in Keras and Torch [42] consistently showed the weakest predictive performance (Tables 1–2), which is likely due to the *p* ≫ *n* regime, typical of microbiome data; the high dimensionality combined with relatively small to moderate sample sizes (*n* ≈ 10^2^ ~ 10^3^), which elevates overfitting risk for high-capacity models whose benefits usually emerge only with much larger samples (e.g., *n >* 10^5^).

### 3.4 Probabilistic Mechanism of PIRF

To illustrate the probabilistic mechanism of PIRF, relative to the other Random Forest extensions: Random Forest [2] and Random Forest (One Cluster) [17], I created a plot displaying the probabilities of microbial features to be selected by each method. To save space, only the plot for the classification task of Obesity [21] is reported in the main text (Fig. 1), while the ones for the remaining tasks are provided in the Supplemental Materials: Inflammation [18] (S1 Fig); Immunotherapy [19] (S2 Fig); T1D [20] (S3 Fig); Cytokine [18] (S4 Fig); Age (Oral) [18] (S5 Fig); and Age (Gut) [21] (S6 Fig).

**Figure 1:**
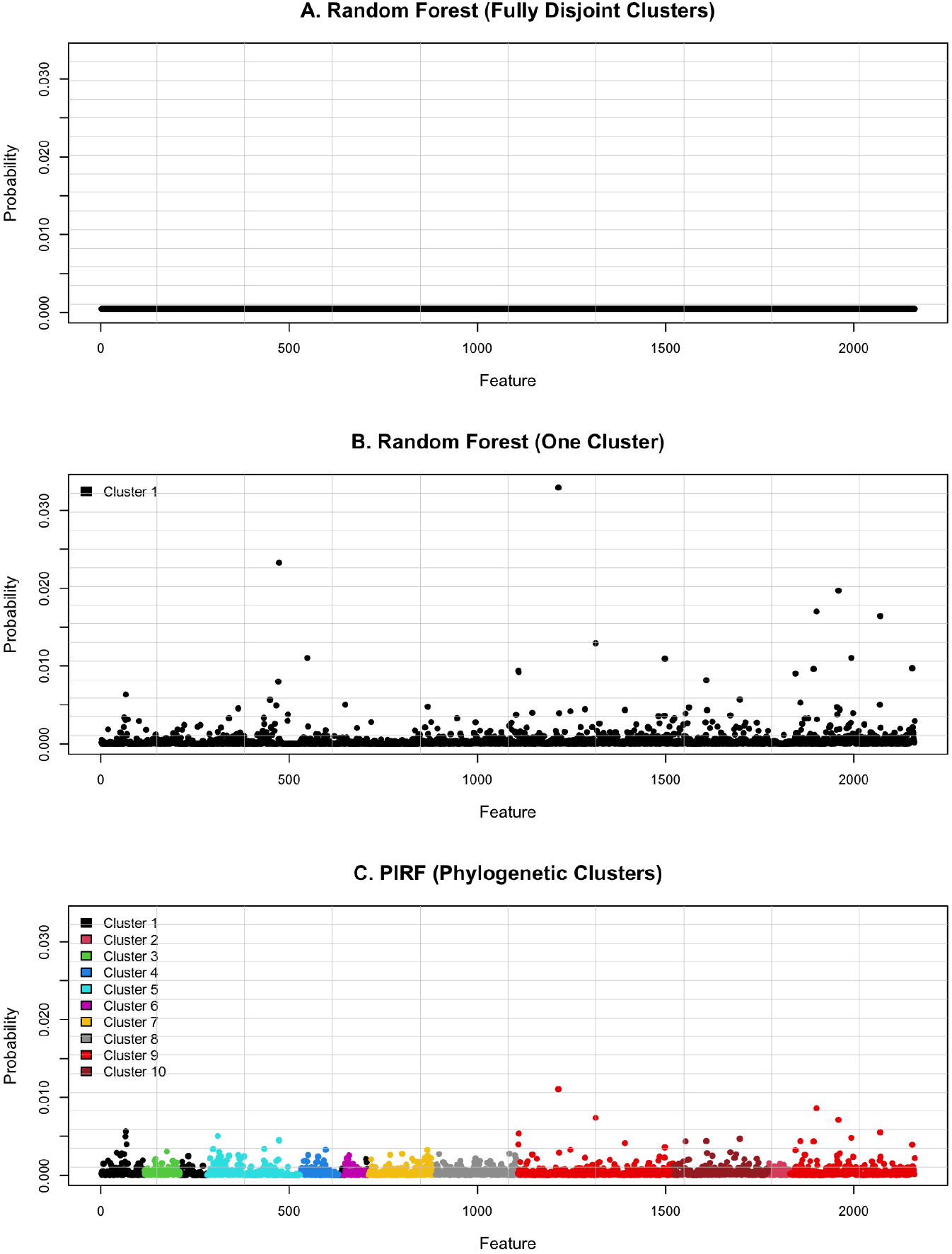
Visual representation of the selection probabilities for the classification task of Obesity using three methods: A. Random Forest (Fully Disjoint Clusters); B. Random Forest (One Cluster); and C. PIRF (Phylogenetic Clusters). Within each method, the probabilities sum to one; as such, Random Forest (Fully Disjoint Clusters) shows the least variability, Random Forest (One Cluster) shows the highest variability, and PIRF (Phylogenetic Clusters) shows an intermediate level of variability.

Note that for each method the probabilities sum to one. The probabilistic mechanisms are described as follows.

- **Random Forest (Fully Disjoint Clusters) [2] (Fig. 1A; S1A - S6A Fig):** Standard Random Forest [2] - which selects features uniformly at random corresponding to the case of fully disjoint clusters (*k* = *p*) - yields the lowest variance in probabilities; and thus, maximizes tree decorrelation, but lacks any feature selection or weighting mechanism.
- **Random Forest (One Cluster) [17] (Fig. 1B; S1B - S6B Fig):** Random Forest based on one community-level cluster - which performs feature selection and weighting within a single cluster (*k* = 1) - yields the highest variance in probabilities; and thus minimizes tree decorrelation, and enforces a globalized feature selection and weighting mechanism.
- **PIRF (Phylogenetic Clusters) (Fig. 1C; S1C - S6C Fig):** Localized Random Forest based on phylogenetic clusters yields intermediate levels of variability in probabilities and tree decorrelation. Importantly, phylogenetic clusters are biologically meaningful groups of microbial features that are evolutionarily and functionally related. This localized strategy enriches functional representations.

To summarize, the standard Random Forest [2] benefits from maximizing tree decorrelation but lacks any mechanism for feature selection and weighting, whereas the Random Forest (One Cluster) [17] emphasizes feature selection and weighting at the expense of substantially reduced tree decorrelation. In contrast, PIRF introduces a localized approach that balances tree decorrelation with feature selection and weighting, while leveraging phylogenetic clusters to diversify functional representations. This balance, coupled with its enriched functional representations, underpins the superior predictive performance of PIRF in microbiome applications.

## 4 Discussion

In this paper, I introduced PIRF, a method designed to improve predictive accuracy in human microbiome studies. The core mechanism of PIRF lies in its localized approach: rather than treating all features as competing globally to be selected or weighted [17], PIRF identifies informative features within each phylogenetic cluster - a localized group of evolutionarily and functionally related microbial features. This strategy enriches functional representations while reducing tree correlation. To achieve it, PIRF partitions the microbial feature space into multiple phylogenetic clusters using phylogenetic tree information. It then computes feature importance scores within each cluster and converts them into cluster-specific probabilities. Finally, these cluster-specific probabilities are integrated across all phylogenetic clusters to derive community-level selection probabilities.

To evaluate predictive performance, I applied PIRF to seven benchmark tasks, comprising four classification problems (Inflammation [18], Immunotherapy [19], T1D [20], and Obesity [21]) and three regression problems (Cytokine [18], Age [Oral] [18], and Age [Gut] [21]). Across these tasks, PIRF achieved the strongest predictive performance - yielding the lowest test error rate and the highest test AUC for classification, and the lowest test RMSE and MAE for regression - outperforming other off-the-shelf tools: Random Forest [2], Random Forest (One Cluster) [17], Gradient Boosting Machine [3], Ridge Regression [36], Lasso [37], Elastic Net [38], and two Deep Neural Network implementations (Keras [39, 40] and Torch [39, 40, 42]).

The software and tutorials are freely available as an R package, `PIRF`, at https://github.com/hk1785/PIRF, whereas many recent methods lack accessible software, limiting their practical utility. PIRF can serve as a useful tool for microbiome-based disease diagnostics and personalized medicine.

## Acknowledgements

HK is the sole author and contributed to every aspect. HK is grateful to the anonymous reviewers for their careful observations and insightful feedback. This work was supported by the National Research Foundation of Korea (NRF) grant funded by the Korean government (MSIT) (2021R1C1C1013861).

## Supplemental Materials

Figures S1–S6 are provided in the supplemental materials.

## References

[1] Breiman L, Friedman JH, Stone CJ. Classification and Regression Trees. CRC Press, Boca Raton, FL, USA„ 1984.

[2] Breiman L. Random Forests. Mach Learn, 45:5–32, 2001.

[3] Friedman JH. Greedy Function Approximation: A Gradient Boosting Machine. Ann Stat, 29:1189–1232, 2001.

[4] Gilbert JA, Blaser MJ, Caporaso JG, Jansson JK, Lynch SV, Knight R. Current Understanding of the Human Microbiome. Nat Med, 24:392–400, 2018.

[5] Faith DP. Conservation evaluation and phylogenetic diversity. Biol Conserv, 61(1):1–10, 1992.

[6] Lozupone C, Knight R. UniFrac: A new phylogenetic method for comparing microbial communities. Appl Environ Microbiol, 71(11):8228–35, 2005.

[7] Anderson MJ. A new method for non-parametric multivariate analysis of variance. Austral Ecol, 26:32–46, 2001.

[8] Zhao N, Chen J, Carroll IM, Ringel-Kulka T, Epstein MP, Zhou H, Zhou JJ, Ringel Y, Li H, Wu MC. Testing in microbiome-profiling studies with MiRKAT, the microbiome regression-based kernel association test. Am J Hum Genet, 96(5):797–807, 2015.

[9] Zhang J, Wei Z, Chen J. A distance-based approach for testing the mediation effect of the human microbiome. Bioinformatics, 34(11):1875–1883, 2018.

[10] Yue Y, Hu YJ. A new approach to testing mediation of the microbiome at both the community and individual taxon levels. Bioinformatics, 38(12):3173–3180, 2022.

[11] Wang Y, Bhattacharya T, Jiang Y, Qin X, Wang Y, Liu Y, Saykin AJ, Chen L. A novel deep learning method for predictive modeling of microbiome data. Brief Bioinform, 22(3):bbaa073, 2021.

[12] Koh H. Subgroup Identification Using Virtual Twins for Human Microbiome Studies. IEEE/ACM Trans Comput Biol Bioinform, 20(6):3800–3808, 2023.

[13] Li B, Wang T, Qian M, Wang S. MKMR: a multi-kernel machine regression model to predict health outcomes using human microbiome data. Brief Bioinform, 24(3):bbad158, 2025.

[14] Xu H, Wang T, Miao Y, Qian M, Yang Y, Wang S. MK-BMC: a Multi-Kernel framework with Boosted distance metrics for Microbiome data for Classification. Bioinformatics, 40(1):btad757, 2024.

[15] Granitto PM, Furlanello C, Biasioli F, Gasperi F. Recursive feature elimination with random forest for PTR-MS analysis of agroindustrial products. Chemometr Intell Lab Syst, 83(2):83–90, 2006.

[16] Darst BF, Malecki KC, Engelman CD. Using recursive feature elimination in random forest to account for correlated variables in high dimensional data. BMC Genet, 19(65):1–6, 2018.

[17] Liu Y, Zhao H. Variable importance-weighted Random Forests. Quant Biol, 5(4):338–351, 2017.

[18] Park B, Koh H, Patatanian M, Reyes-Caballero H, Zhao N, Meinert J, Holbrook JT, Leinbach LI, Biswal S. The mediating roles of the oral microbiome in saliva and subgingival sites between e-cigarette smoking and gingival inflammation. BMC Microbiol, 23(35), 2023.

[19] Limeta A, Ji B, Levin M, Gatto F, Nielsen J. Meta-analysis of the gut microbiota in predicting response to cancer immunotherapy in metastatic melanoma. JCI Insight, 5(23):e140940, 2020.

[20] Zhang XS, Li J, Krautkramer KA, et al. Antibiotic-induced acceleration of type 1 diabetes alters maturation of innate intestinal immunity. eLife, 7(e37816):1–37, 2018.

[21] McDonald D, Hyde E, Debelius JW, et al. American Gut Consortium; Knight R. American Gut: an Open Platform for Citizen Science Microbiome Research. mSystems, 3(3):e00031–18, 2018.

[22] Wright MN, Ziegler A. ranger: A Fast Implementation of Random Forests for High Dimensional Data in C++ and R. J Stat Softw, 77(1):1–17, 2017.

[23] Sneath PHA, Sokal RR, Freeman WH. Numerical taxonomy: The principles and practice of numerical classification. Syst Zool, 24:263–8, 1975.

[24] Reynolds AP, Richards G, Iglesia B, Rayward-Smith VJ. Clustering rules: A comparison of partitioning and hierarchical clustering algorithms. J Math Model Algorithms, 5(4):475–504, 2006.

[25] Koh H, Zhao N. A powerful microbial group association test based on the higher criticism analysis for sparse microbial association signals. Microbiome, 8(63), 2020.

[26] Rousseeuw PJ. Silhouettes: A graphical aid to the interpretation and validation of cluster analysis. J Comput Appl Math, 20:53–65, 1987.

[27] Altmann A, Tolosi L, Sander O, Lengauer T. Permutation importance: a corrected feature importance measure. Bioinformatics, 26(10):1340–1347, 2010.

[28] Louppe G, Wehenkel L, Sutera A, Geurts P. Understanding variable importances in forests of randomized trees. In Advances in Neural Information Processing Systems (NIPS), volume 26, 2013.

[29] Ling W, Lu J, Zhao N, Lulla A, Plantinga AM, Fu W, Zhang A, Liu H, Song H, Li Z, et al. Batch effects removal for microbiome data via conditional quantile regression. Nat Commun, 13(5418), 2022.

[30] Matson V, Fessler J, Bao R, Chongsuwat T, Zha Y, Alegre ML, Luke JJ, Gajewski TF. The commensal microbiome is associated with anti–PD-1 efficacy in metastatic melanoma patients. Science, 359(6371):104–8, 2018.

[31] Frankel AE, Coughlin LA, Kim J, Froehlich TW, Xie Y, Frenkel EP, Koh AY. Metagenomic shotgun sequencing and unbiased metabolomic profiling identify specific human gut microbiota and metabolites associated with immune checkpoint therapy efficacy in melanoma patients. Neoplasia, 19(10):848–55, 2017.

[32] Gopalakrishnan V, Spencer CN, Nezi L, Reuben A, Andrews MC, Karpinets T, Prieto PA, Vicente D, Hoffman K, Wei SC, et al. Gut microbiome modulates response to anti–PD-1 immunotherapy in melanoma patients. Science, 359(6271):97–103, 2018.

[33] Routy B, Le Chatelier E, Derosa L, Duong CP, Alou MT, Daillere R, Fluckiger A, Messaoudene M, Rauber C, Roberti MP, et al. Gut microbiome influences efficacy of PD-1–based immunotherapy against epithelial tumors. Science, 359(6371):91–7, 2018.

[34] Peters BA, Wilson M, Moran U, Pavlick A, Izsak A, Wechter T, Weber JS, Osman I, Ahn J. Relating the gut metagenome and metatranscriptome to immunotherapy responses in melanoma patients. Genome Med, 11(61), 2019.

[35] Chen T, Guestrin C. Xgboost: A scalable tree boosting system. In Proceedings of the 22nd ACM SIGKDD International Conference on Knowledge Discovery and Data Mining (KDD), pages 785–794, 2016.

[36] Hoerl AE, Kennard RW. Ridge regression: Biased estimation for nonorthogonal problems. Technometrics, 12(1):55–67, 1970.

[37] Tibshirani R. Regression shrinkage and selection via the lasso. J R Stat Soc B, 58(1):267–288, 1996.

[38] Zou H, Hastie T. Regularization and variable selection via the elastic net. J R Stat Soc B, 67(2):301–320, 2005.

[39] Hinton GE, Osindero S, Teh YW. A fast learning algorithm for deep belief nets. Neural Comput, 18(7):1527–1554, 2006.

[40] LeCun Y, Bengio Y, Hinton G. Deep learning. Nature, 521(7553):436–444, 2015.

[41] Kingma DP, Ba J. Adam: A Method for Stochastic Optimization. In Proceedings of the 3rd International Conference on Learning Representations (ICLR), 2015.

[42] Paszke A, Gross S, Massa F, Lerer A, Bradbury J, Chanan G, Killeen T, Lin Z, Gimelshein N, Antiga L, Desmaison A, Kopf A, Yang E, DeVito Z, Raison M, Tejani A, Chilamkurthy S, Steiner B, Fang L, Bai J, Chintala S. PyTorch: An Imperative Style, High-Performance Deep Learning Library. In Advances in Neural Information Processing Systems (NeurIPS), pages 8024–8035, 2019.

