## Supplementary Materials for "Phylogeny-Informed Random Forests for Human Microbiome Studies"

**S1 Fig.** Visual representation of the selection probabilities for the classification task of Inflammation using three methods: A. Random Forest (Fully Disjoint Clusters); B. Random Forest (One Cluster); and C. PIRF (Phylogenetic Clusters). Within each method, the probabilities sum to one; as such, Random Forest (Fully Disjoint Clusters) shows the least variability, Random Forest (One Cluster) shows the highest variability, and PIRF (Phylogenetic Clusters) shows an intermediate level of variability.

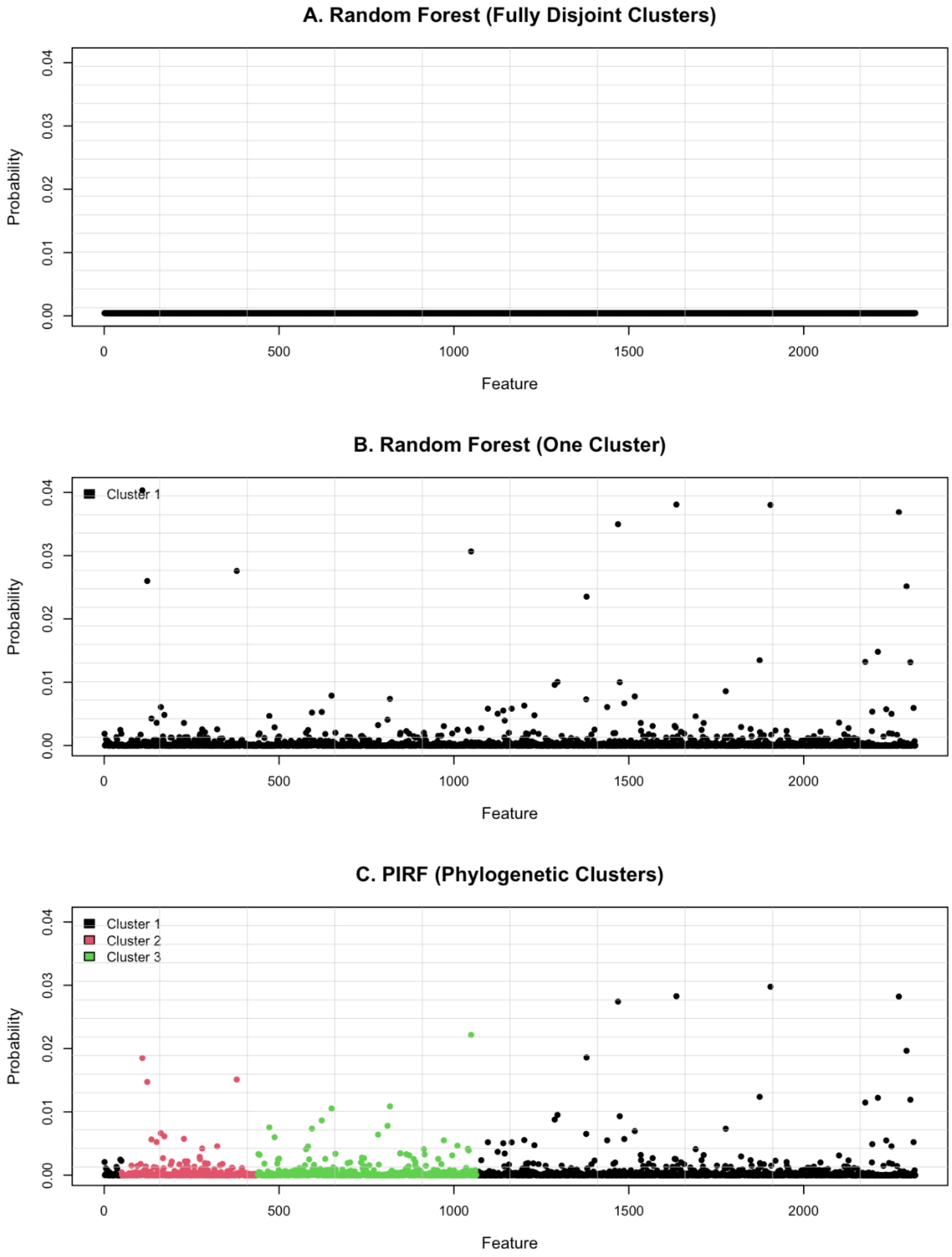

**S2 Fig.** Visual representation of the selection probabilities for the classification task of Immunotherapy using three methods: A. Random Forest (Fully Disjoint Clusters); B. Random Forest (One Cluster); and C. PIRF (Phylogenetic Clusters). Within each method, the probabilities sum to one; as such, Random Forest (Fully Disjoint Clusters) shows the least variability, Random Forest (One Cluster) shows the highest variability, and PIRF (Phylogenetic Clusters) shows an intermediate level of variability.

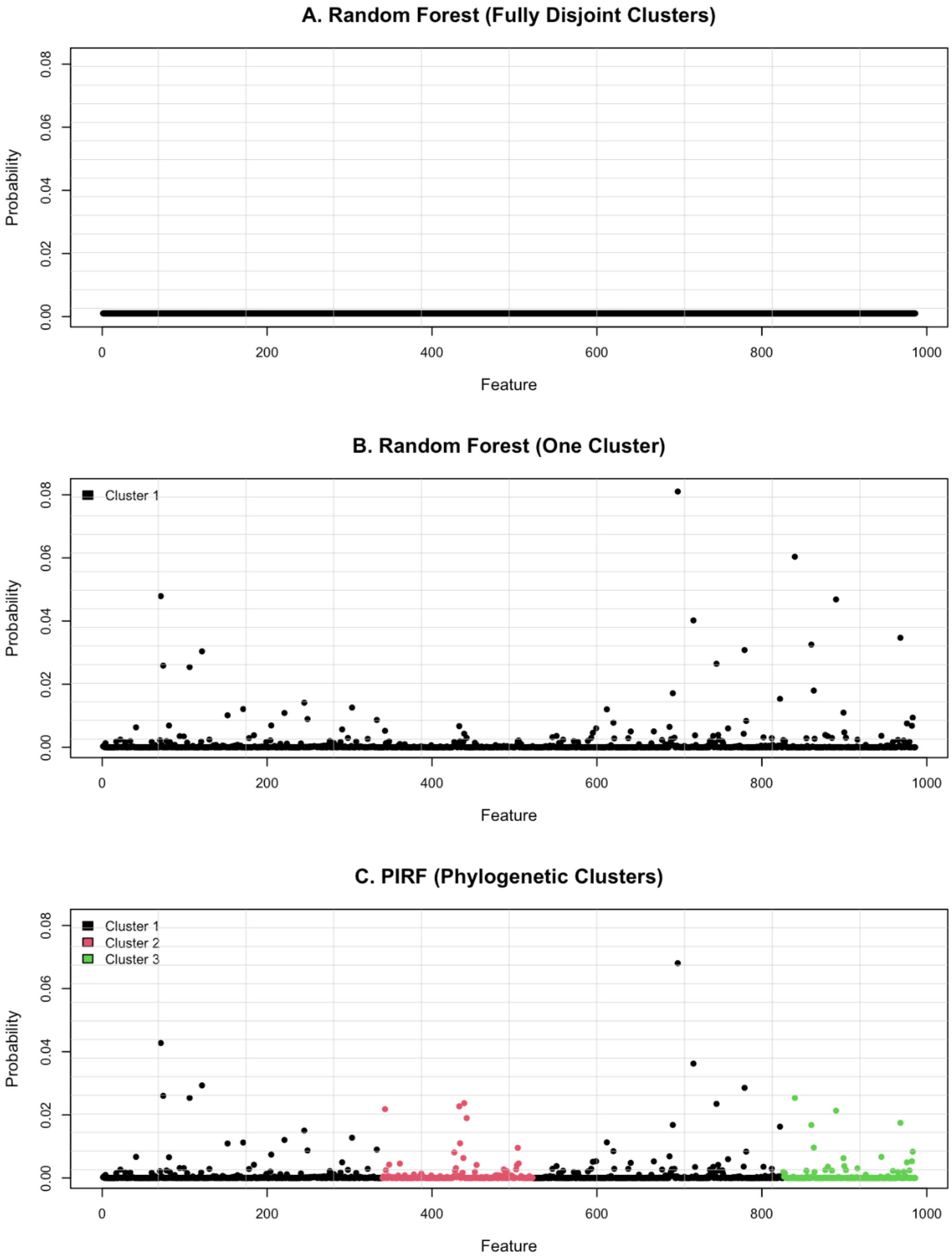

**S3 Fig.** Visual representation of the selection probabilities for the classification task of T1D using three methods: A. Random Forest (Fully Disjoint Clusters); B. Random Forest (One Cluster); and C. PIRF (Phylogenetic Clusters). Within each method, the probabilities sum to one; as such, Random Forest (Fully Disjoint Clusters) shows the least variability, Random Forest (One Cluster) shows the highest variability, and PIRF (Phylogenetic Clusters) shows an intermediate level of variability.

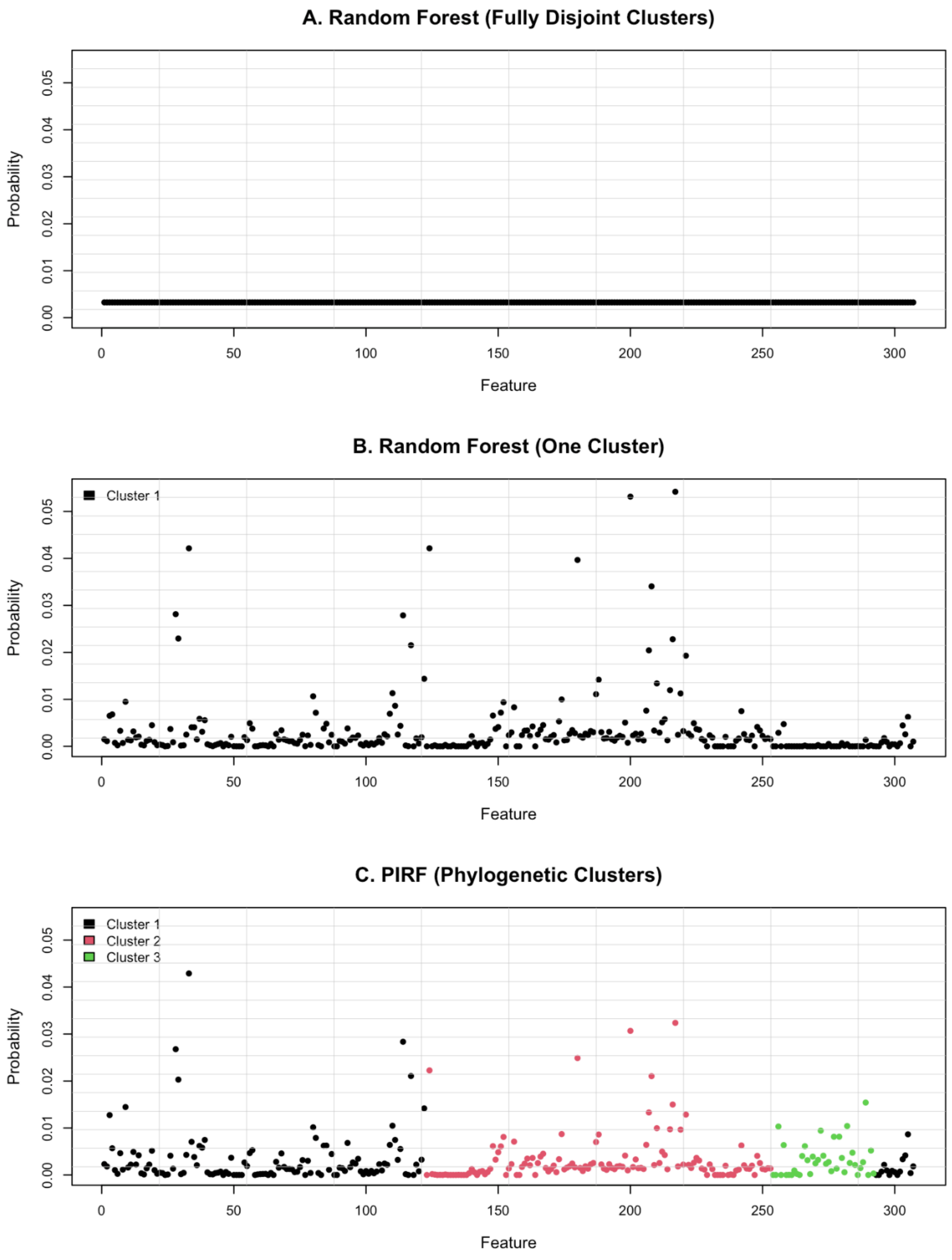

**S4 Fig.** Visual representation of the selection probabilities for the regression task of Cytokine using three methods: A. Random Forest (Fully Disjoint Clusters); B. Random Forest (One Cluster); and C. PIRF (Phylogenetic Clusters). Within each method, the probabilities sum to one; as such, Random Forest (Fully Disjoint Clusters) shows the least variability, Random Forest (One Cluster) shows the highest variability, and PIRF (Phylogenetic Clusters) shows an intermediate level of variability.

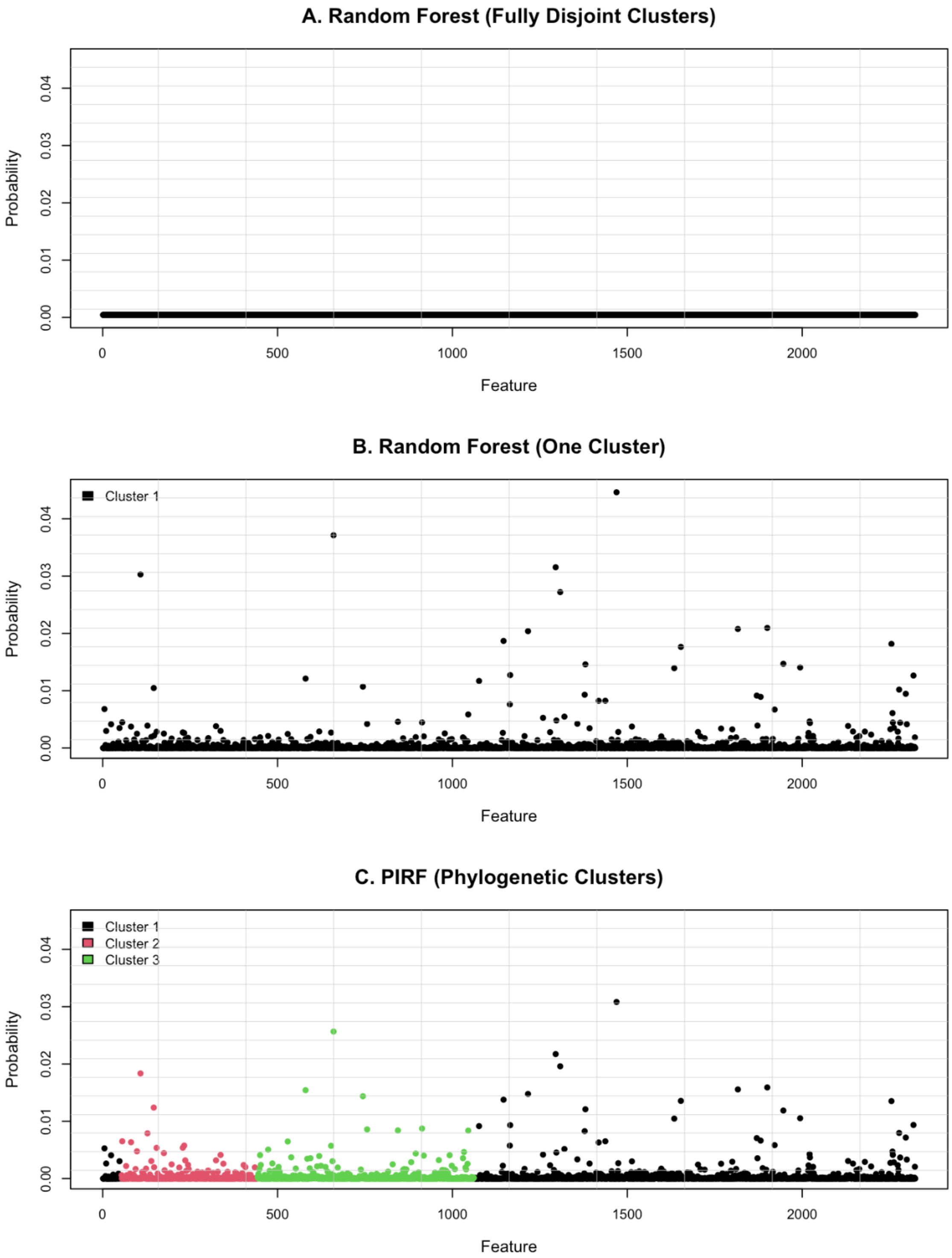

**S5 Fig.** Visual representation of the selection probabilities for the regression task of Age (Oral) using three methods: A. Random Forest (Fully Disjoint Clusters); B. Random Forest (One Cluster); and C. PIRF (Phylogenetic Clusters). Within each method, the probabilities sum to one; as such, Random Forest (Fully Disjoint Clusters) shows the least variability, Random Forest (One Cluster) shows the highest variability, and PIRF (Phylogenetic Clusters) shows an intermediate level of variability.

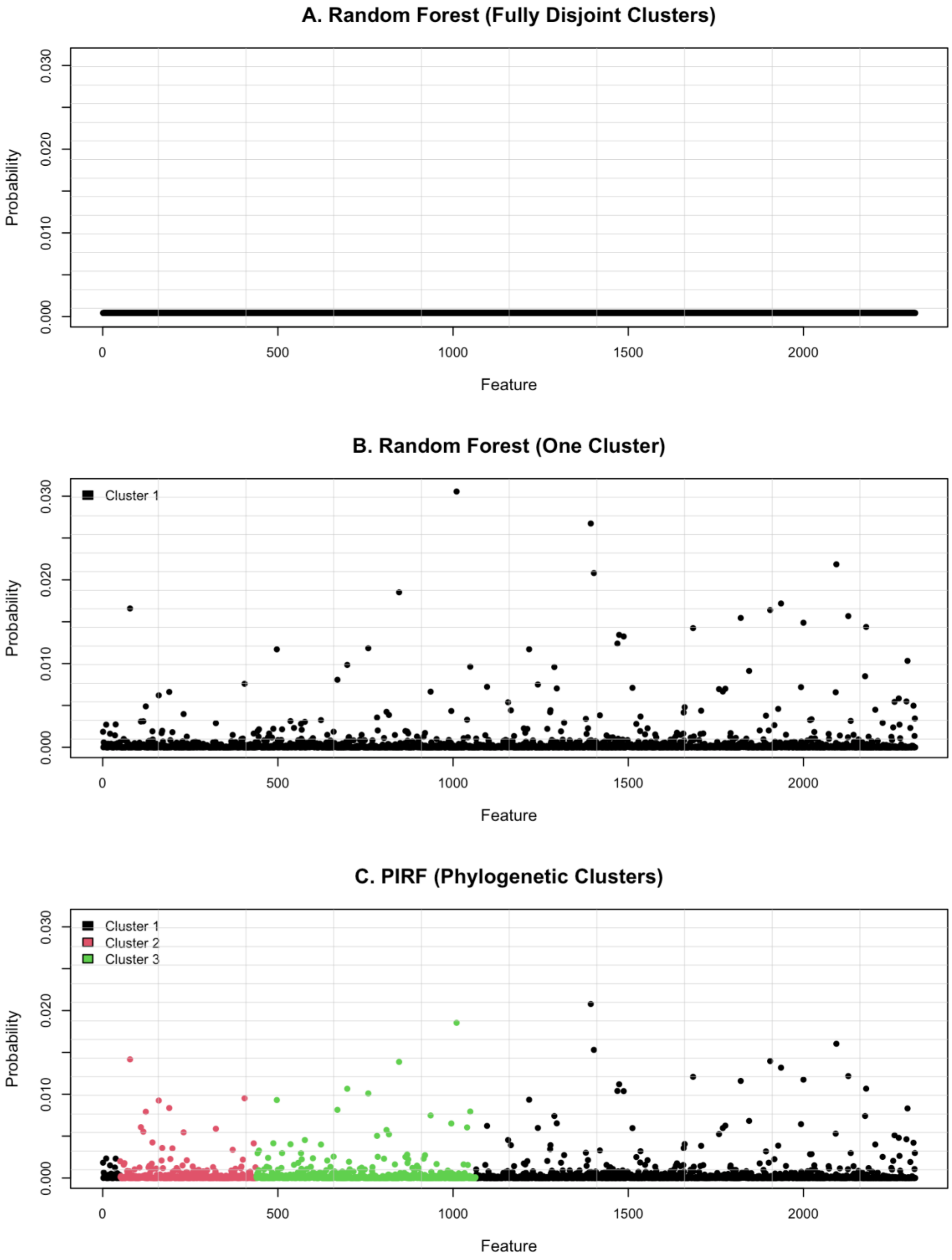

**S6 Fig.** Visual representation of the selection probabilities for the regression task of Age (Gut) using three methods: A. Random Forest (Fully Disjoint Clusters); B. Random Forest (One Cluster); and C. PIRF (Phylogenetic Clusters). Within each method, the probabilities sum to one; as such, Random Forest (Fully Disjoint Clusters) shows the least variability, Random Forest (One Cluster) shows the highest variability, and PIRF (Phylogenetic Clusters) shows an intermediate level of variability.

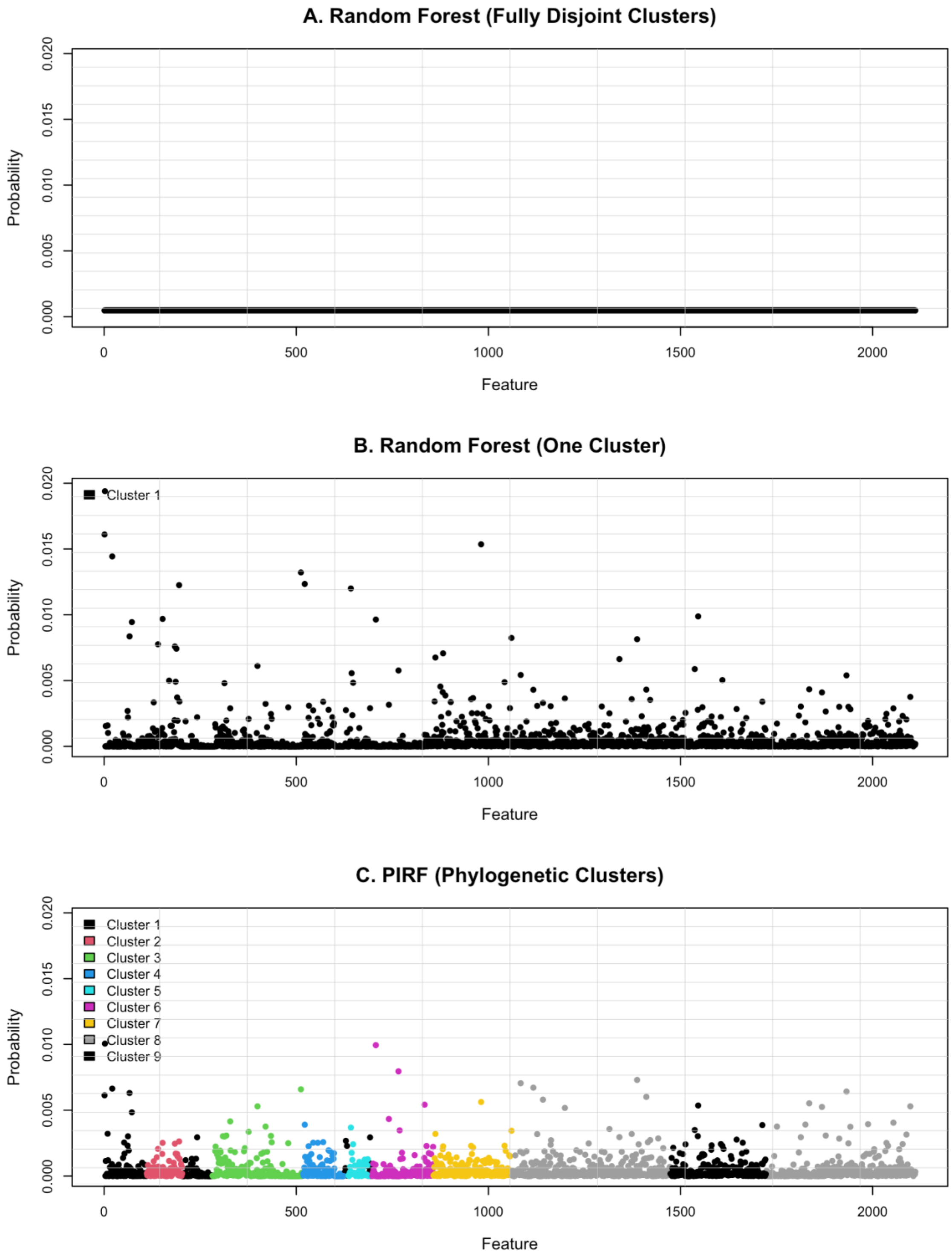
